# Prospective Isolation of Chondroprogenitors from Human iPSCs Based on Cell Surface Markers Identified using a CRISPR-Cas9-Generated Reporter

**DOI:** 10.1101/675983

**Authors:** Amanda Dicks, Chia-Lung Wu, Nancy Steward, Shaunak S. Adkar, Charles A. Gersbach, Farshid Guilak

## Abstract

Articular cartilage shows little or no capacity for intrinsic repair, generating a critical need for regenerative therapies for joint injuries and diseases such as osteoarthritis. Human induced pluripotent stem cells (hiPSCs) offer a promising cell source for cartilage tissue engineering and *in vitro* human disease modeling; however, heterogeneity and off-target differentiation remain a challenge. We used a CRISPR-Cas9-edited *COL2A1-GFP* knock-in reporter hiPSC line, coupled with a surface marker screen, to identify a novel chondroprogenitor population expressing CD146, CD166, and PDGFRβ, but not CD45. Under chondrogenic culture conditions, these triple positive chondroprogenitor cells demonstrated decreased heterogeneity as measured by single cell RNA sequencing, as well as more robust and homogenous matrix production with significantly higher chondrogenic gene expression. Overall, this study has identified a unique hiPSC-derived subpopulation of chondroprogenitors that are CD146^+^/CD166^+^/PDGFRβ^+^/CD45^-^ and exhibit high chondrogenic potential, providing a purified cell source for cartilage tissue engineering or disease modeling studies.

## INTRODUCTION

Articular cartilage is the load-bearing tissue that lines the ends of long bones in diarthrodial joints, serving to resist compression and provide a nearly frictionless surface during joint loading and movement (Guilak, 2011; Mansour, 2009). The extracellular matrix of cartilage is comprised primarily of type II collagen and proteoglycans, which are synthesized by the main residing cell type, chondrocytes (Fox et al., 2009; Lin et al., 2006). However, because it is aneural and avascular, cartilage shows little or no capacity for intrinsic repair (Fox et al., 2009). Traumatic injury and a chronic inflammatory state lead to irreversible degeneration of the tissue, driving diseases such as osteoarthritis (OA) (Berenbaum et al., 2016; Lieberthal et al., 2015). Current treatments only target disease symptoms, creating a great demand for tissue-engineered cartilage as a system for disease modeling, drug testing, and tissue replacement.

Human induced pluripotent stem cells (hiPSCs) offer a promising source for cartilage tissue engineering and *in vitro* disease modeling (Musunuru, 2013) as they have virtually unlimited expansion capacity, can be genetically modified, and avoid many of the ethical considerations associated with embryonic stem cells (Takahashi et al., 2007; Yumlu et al., 2017). Despite reports of several chondrogenic differentiation protocols for pluripotent stem cells (Craft et al., 2015; Lee et al., 2015; Lian et al., 2010; Nejadnik et al., 2015; Suchorska et al., 2017; Yamashita et al., 2015), incomplete differentiation and cell heterogeneity remain major obstacles for iPSC chondrogenesis (Cahan and Daley, 2013; Yoshida and Yamanaka, 2010). This challenge has been addressed in other stem and progenitor cell types by prospectively isolating cells that exhibit chondrogenic lineage commitment using surface marker expression. For example, previous studies have identified chondroprogenitors within adult articular cartilage that can be isolated using fibronectin adhesion assays since progenitors express integrins α5 and β1 (Vinod et al., 2018; Williams et al., 2010). Additionally, mesenchymal progenitor cells, which express CD105, CD166, and CD146, have been reported to have a high chondrogenic potential (Alsalameh et al., 2004; Su et al., 2015; Vinod et al., 2018). Adult multipotent cells, such as bone marrow mesenchymal stem cells (MSCs) or adipose stem cells (ASCs), exhibit chondrogenic potential and have been used extensively for cartilage tissue engineering. They are often characterized by a range of cell surface marker expression, including CD105, CD73, CD90, CD271, CD146, Stro-1, and SSEA-4 (Lv et al., 2014). In an effort to identify a more developmentally-relevant progenitor population, self-renewing human skeletal stem cells characterized by CD164^+^, CD73^-^, and CD146^-^ showed chondrogenic differentiation when implanted in a mouse renal capsule (Chan et al., 2018). In another study, limb bud cells expressing CD73 and BMPR1β while having low to no expression of CD166, CD146, and CD44 were proposed to be the earliest cartilage committed cells (prechondrocytes) in human embryonic development (Wu et al., 2013). However, surface markers characteristic of hiPSC-derived chondroprogenitors or chondrocytes remain to be identified.

Previously, our lab used green fluorescent protein (GFP) reporter systems to track the expression of collagen type II alpha 1 chain (*COL2A1*) in mouse (Diekman et al., 2012) and human (Adkar et al., 2018) iPSCs, allowing for the prospective isolation and purification of *COL2A1*-GFP+ chondrogenic cells during the differentiation process. Despite the fact that this approach significantly enhanced iPSC chondrogenesis (Adkar et al., 2018), genome editing is required to create a reporter line, hindering potential clinical translation. In this regard, the identification of cell surface markers that are directly representative of this *COL2A1*-positive population could greatly enhance the prospective isolation and purification of chondroprogenitors, without requiring genetic modifications to the cell line.

In this study, we used a *COL2A1*-GFP knock-in reporter hiPSC line as a tool to identify cell surface markers that are highly co-expressed with *COL2A1* to test the hypothesis that this sub-population of chondroprogenitor cells will show increased purity and chondrogenic capacity. Single-cell RNA sequencing (scRNA-seq) was then used to investigate the gene expression profile of this population and to identify subsets within it. Matrix production, cell morphology, and gene expression were measured to evaluate chondrogenic ability of unsorted and sorted chondroprogenitor cells. This chondroprogenitor population appears to represent an intermediate step in the developmental pathway of hiPSC differentiation into chondrocytes. The identification of a purified population of chondroprogenitor cells will enhance the efficiency of hiPSC-chondrogenic differentiation for use in tissue engineering, *in vitro* disease modeling, and drug testing.

## RESULTS

### *COL2A1*-positive chondroprogenitor cells express PDGFRβ, CD146, and CD166

*COL2A1*-GFP reporter hiPSCs were differentiated into chondroprogenitor cells along the mesodermal lineage as previously described (Adkar et al., 2018). After differentiation, the cells were labeled for surface markers commonly associated with MSCs and/or chondroprogenitors in the developing limb bud (Chan et al., 2018; Lv et al., 2014; Wu et al., 2013). Flow cytometric analysis showed that, on average, 4.27% of the chondroprogenitor population expressed *COL2A1* based on GFP expression (**Figure S1A**). Of the total population, less than 1% also expressed CD271, CD105, CD73, and BMPR1β. The chondroprogenitor cells had the highest expression of CD271 of the four markers, but it was not co-expressed with *COL2A1* (**Figure S1C)**. Interestingly, PDGFRβ, CD146, and CD166 were co-expressed with *COL2A1* at the highest frequencies as 2.32%, 2.17%, and 1.32% of the total population, respectively **(Figure S1B)**. Therefore, the presence of these three surface markers, and the absence of the hematopoietic marker CD45, were used to sort for chondroprogenitor cells (**Figure 1A)**.

**Figure 1.**
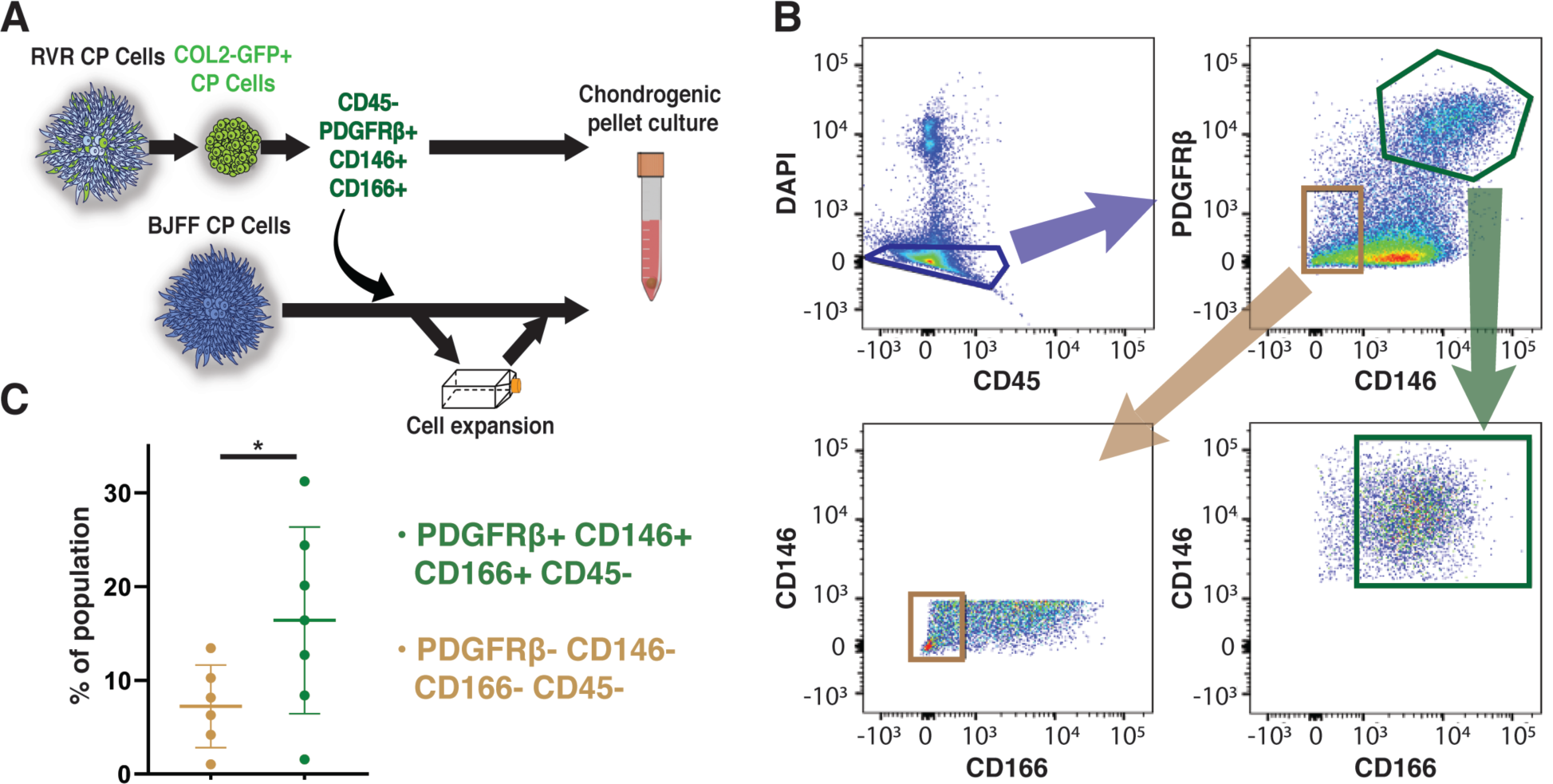
Sorting strategy to identify progenitors with robust chondrogenic potential from heterogenous chondroprogenitor (CP) cells. (A) A schematic representing the experimental design. The RVR cell line with the *COL2A1*-GFP reporter was differentiated into chondroprogenitor cells. Surface marker analysis indicated that PDGFRβ, CD146, and CD166 expression were highly co-expressed with *COL2A1* but not CD45. (B) Cells expressing these desired markers were sorted from wildtype BJFF chondroprogenitor cells. To evaluate chondrogenic potential of the sorted cells, pellets from the sorted cells were either made immediately post-sorting or formed after *in* vitro expansion. (C) A higher percentage of the total cell population (∼16.5%) were triple positive for the desired markers compared to the population not expressing any of these markers. * p < 0.05. Data represented as mean ± SEM. n = 6-7 independent experiments, with 3 technical replicates per experiment. See also Figure S1.

### Chondroprogenitor cells are triple positive for PDGFRβ, CD146, and CD166

The BJFF hiPSC line (wildtype without genome editing) was differentiated into chondroprogenitor cells accordingly. Cells either directly underwent chondrogenic pellet culture, were expanded, were saved for scRNA-seq, or were labeled for the surface markers of interest (**Figure 1A**). Fluorescent activated cell sorting (FACS) was used to sort live chondroprogenitor cells negative for CD45 and positively expressing PDGFRβ and CD146, followed by expression of CD166 (**Figure 1B)**. Cells not expressing any of these surface markers were also analyzed as a negative control. Approximately 16.5% of the total chondroprogenitor cell population was triple positive for PDGFRβ, CD146, and CD166, which was significantly higher than the percentage of the cells (7.2% of total) that were triple-negative for these markers (**Figure 1C**). Sorted cells were collected and either pelleted for chondrogenesis, expanded or saved for scRNA-seq, as described in **Figure 1A**.

### scRNA-seq reveals that unsorted chondroprogenitor cells contained diverse cell populations

We next used scRNA-seq to explore the cell diversity and genetic profiles of unsorted chondroprogenitor cells. At least 9 distinct cell populations (cell clusters) were observed in unsorted chondroprogenitor cells (**Figure 2A**). Among these populations, 5 of them were enriched for a variety of neural cell markers such as *SOX2, OTX1, NES*, and *PAX6* (**Figure 2B**), likely representing populations of the neurogenic lineage. Furthermore, we found that 3 cell populations exhibited high expression levels of several mesenchyme markers including *PRRX1, COL1A1, COL5A1*, and *COL6A1*, while only a small cell population (2.3% of total) expressed chondrogenic markers such as *SOX9, COL2A1, IGFBP5*, and *NKX3-2* (**Figure 2C**). To understand biological function of each population, we then perform Gene Ontology (GO) enrichment analysis of the gene sets representing each cell cluster (**Figure 2D and S2A)**. We observed that *SOX2/TTR/NES+* cells were enriched for genes of forebrain development, while cells expressing *SOX9* and *COL2A1* demonstrated gene sets enriched for protein translation and skeletal system development. Interestingly, *SOX2/SOX3/TOP2A+* and *PRRX1/HIST1H4C/CDK1+* cell populations, found in neurogenic and mesenchymal lineages, respectively, were both enriched for cell cycle genes **(Figure 2D**). We subsequently assigned each individual cell a cell cycle phase (i.e., G1, S, or G2M) based on the expression of cell cycle genes. We found that 75% of *SOX9/COL2A1* were in quiescent, non-proliferating G1 state.

**Figure 2.**
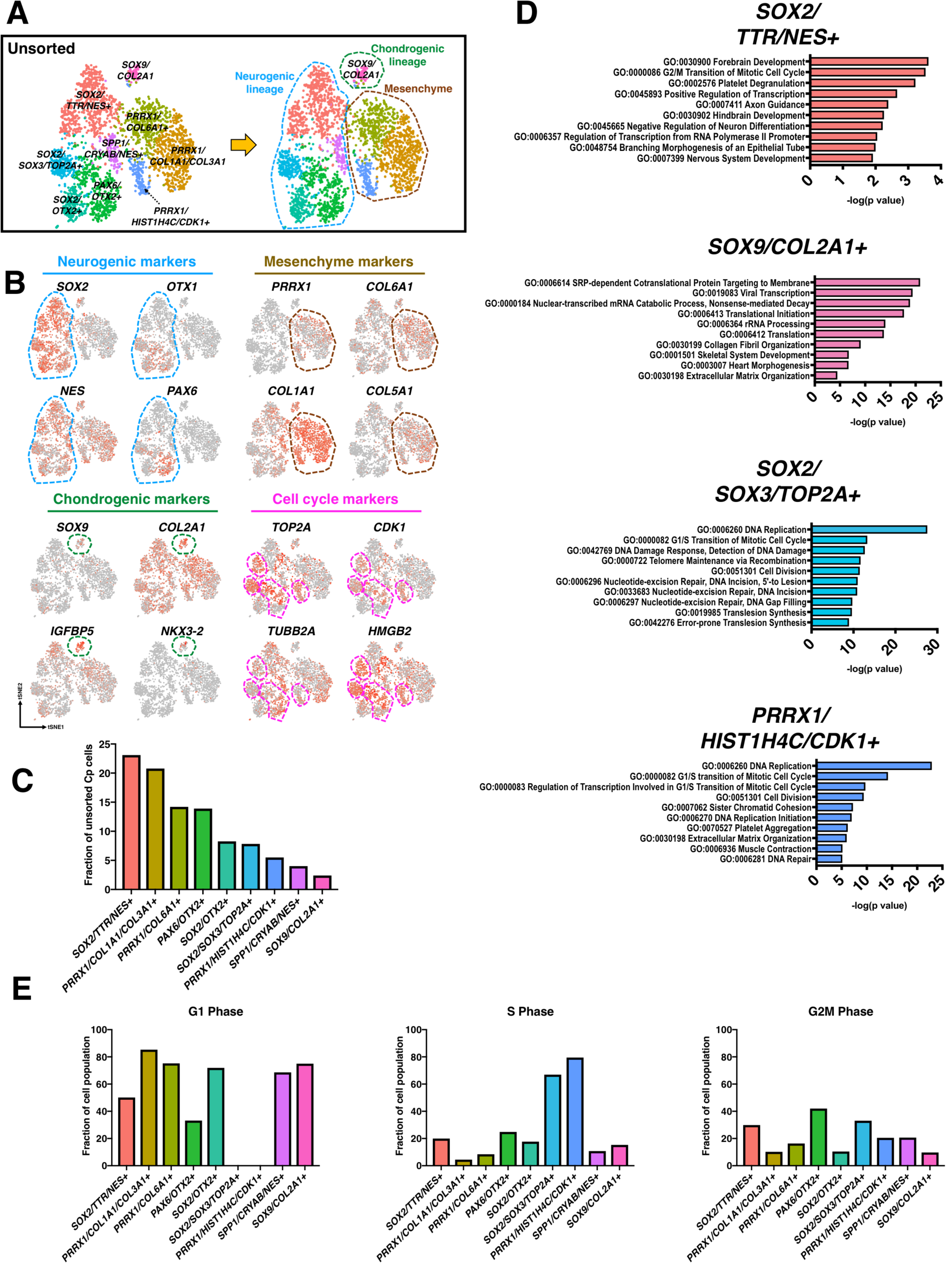
Cell populations and GO enrichment analysis of unsorted chondroprogenitor cells. (A) scRNA-seq identified unsorted chondroprogenitor cells contained at least 9 populations, which could be further categorized into 3 broad classes: neurogenic cells (blue dashed circle), chondrogenic cells (green dashed circle), and mesenchyme (brown dashed circle). (B) Expression of signature genes of each cell lineage. (C) Percentage of total unsorted chondroprogenitor cells in each unique cell population. More than 20% of the unsorted chondroprogenitors were *SOX2/TTR/NES+* neurogenic cells, while only small number of unsorted cells expressed *SOX9* and *COL2A1*. (D) GO terms analysis (biological process) of each unique population. (E) Cell cycle analysis indicates *SOX2/SOX3/TOP2A*+ cells (from neurogenic lineage) and *PRRX1/HIST1H4C/CDK1*+ (from mesenchyme) were proliferative cells. See also Figure S2 and S3.

### scRNA-seq reveals that sorting enriched *SOX9/COL2A1+* cells

scRNA-seq of sorted chondroprogenitor cells indicated that there were at least 6 cell populations consisting of PDGFRβ^+^/CD146^+^/CD166^+^ cells (**Figure 3A)**. Surprisingly, there was still a small percentage of cells (4% of total sorted cells) expressing *SOX2* and *NES*, despite the stringent sorting regime (**Figures 3B and 3C**). We also observed that *SOX2*/*NES+* cells exhibited high expression of CD47, an integrin-associated protein (Brown and Frazier, 2001) (**Figure S2B**). Nevertheless, sorting still significantly enriched cells positive for *SOX9* and *COL2A1* by > 11-fold (27% of total sorted cells). Additionally, we also observed that gene expression levels of the sorting makers were enriched in the sorted cells expressing *SOX9* and *COL2A1* **(Figure 3D)**. Similarly, there was enrichment of some previously reported pro-chondrogenic markers (Alsalameh et al., 2004; Vinod et al., 2018; Williams et al., 2010; Wu et al., 2013) in the sorted chondroprogenitor population; specifically *ITGA5* and *ENG* (CD105) (**Figure S3**). Skeletal system development, as expected, emerged as a significant GO term in *SOX9/COL2A1+* cells, while *HMGB2/TOP2A+* and *LGALS1/PTTG1+* cells were enriched in gene sets of cell division (**Figure 3E and S2C)**. Similar to unsorted cells, the majority of S*OX9/COL2A1+* cells (> 80%) were in quiescent G1 status of cell cycle (**Figure 3F)**.

**Figure 3.**
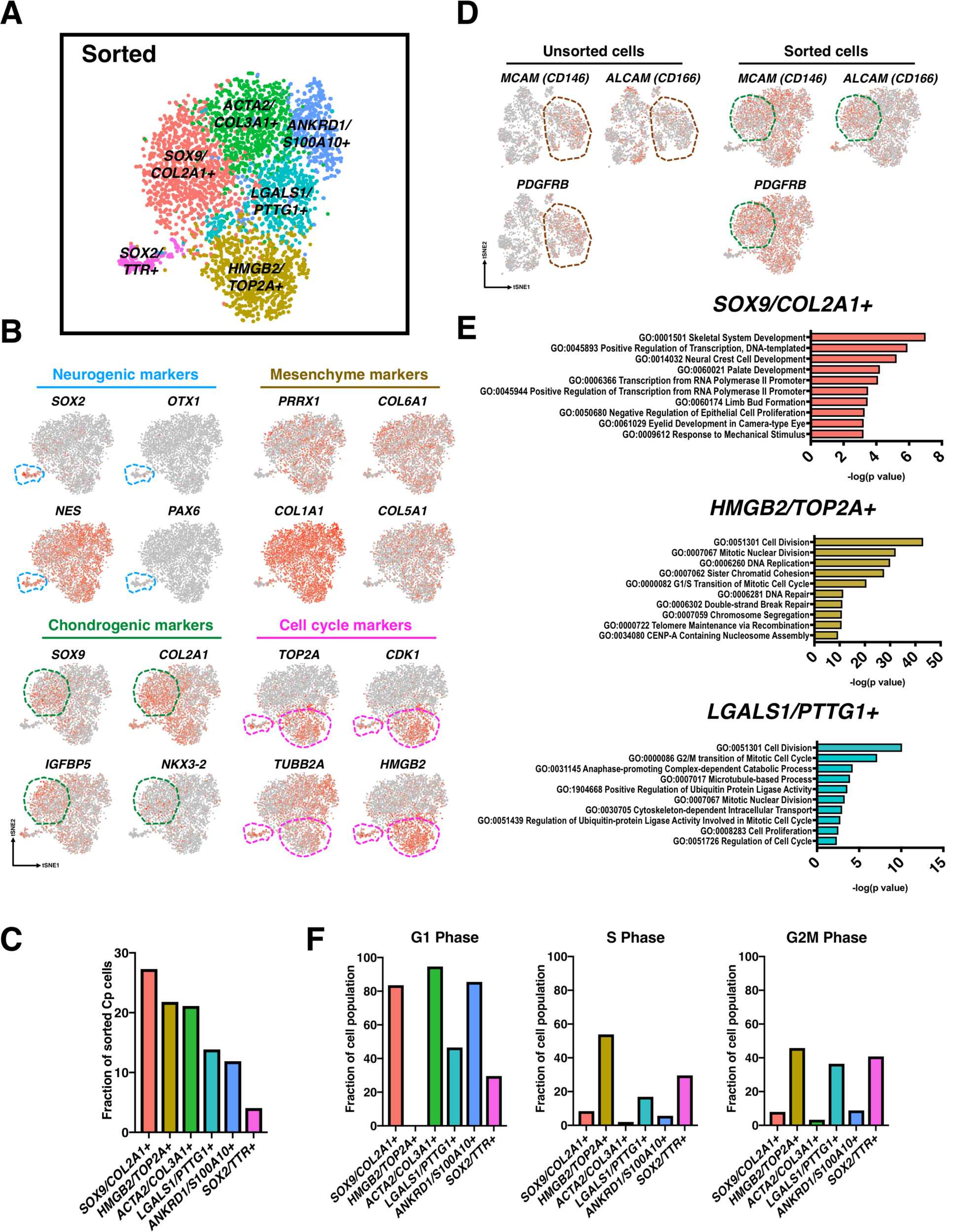
Cell populations and GO enrichment analysis of sorted chondroprogenitor cells. (A) scRNA-seq identified PDGFRβ^+^/CD146^+^/CD166^+^ cells contained at least 6 populations. (B) The sorted cells were enriched for mesenchymal and chondrogenic genes. (C) Percentage of total sorted chondroprogenitor cells in each unique cell population. 27% of the sorted were *SOX9/COL2A1.* Interestingly, a small percentage of cells (4% of total sorted cells) expressing *SOX2* and *NES* was still observed. (D) PDGFRβ^+^/CD146^+^/CD166^+^ sorted cells may belong to mesenchymal population (brown dashed circle) in unsorted cells. Green dashed circle indicates the population that was positive for *SOX9* and *COL2A1.* (E) GO terms analysis (biological process) showing skeletal system development was highlighted in *SOX9/COL2A1+* cells, while *HMGB2/TOP2A+* and *LGALS1/PTTG1+* cells were enriched in gene sets of cell division. (F) Cell cycle analysis indicating that most S*OX9/COL2A1+* cells were in quiescent G1 status of cell cycle. See also Figure S2 and S3.

### Canonical correlation analysis (CCA) analysis demonstrates high enrichment of proliferative and mesenchymal genes in sorted chondroprogenitor cells

Five major conserved populations were identified after CCA alignment of the sorted and unsorted chondroprogenitor cells (**Figure 4A**). Among these populations, *HIST1H4C+* cells accounted for the major conserved population, while the *IGFBP5/COL2A1+* cluster was the smallest. We next performed differentially expression gene (DEG) analysis to explore how sorting enriches or depletes the levels of gene expression within each individual population **(Figure 4B)**. Within the *IGFBP5/COL2A1+* population, sorted cells exhibited significantly up-regulated expression of several mesenchymal genes including *TPM1, TAGLN* and *TMSB10* (indicated by brown circle), which have been suggested to be essential in chondrogenesis (Molnar et al., 2007; Zhang et al., 2018). Furthermore, within *IGFBP5/COL2A1+* population, sorted cells demonstrated significantly down-regulated expression of *IGFBP5* (indicated by blue circle), an important transcription factor inducing chondroprogenitor cells into chondrogenic lineage (Clemmons, 1993).

**Figure 4.**
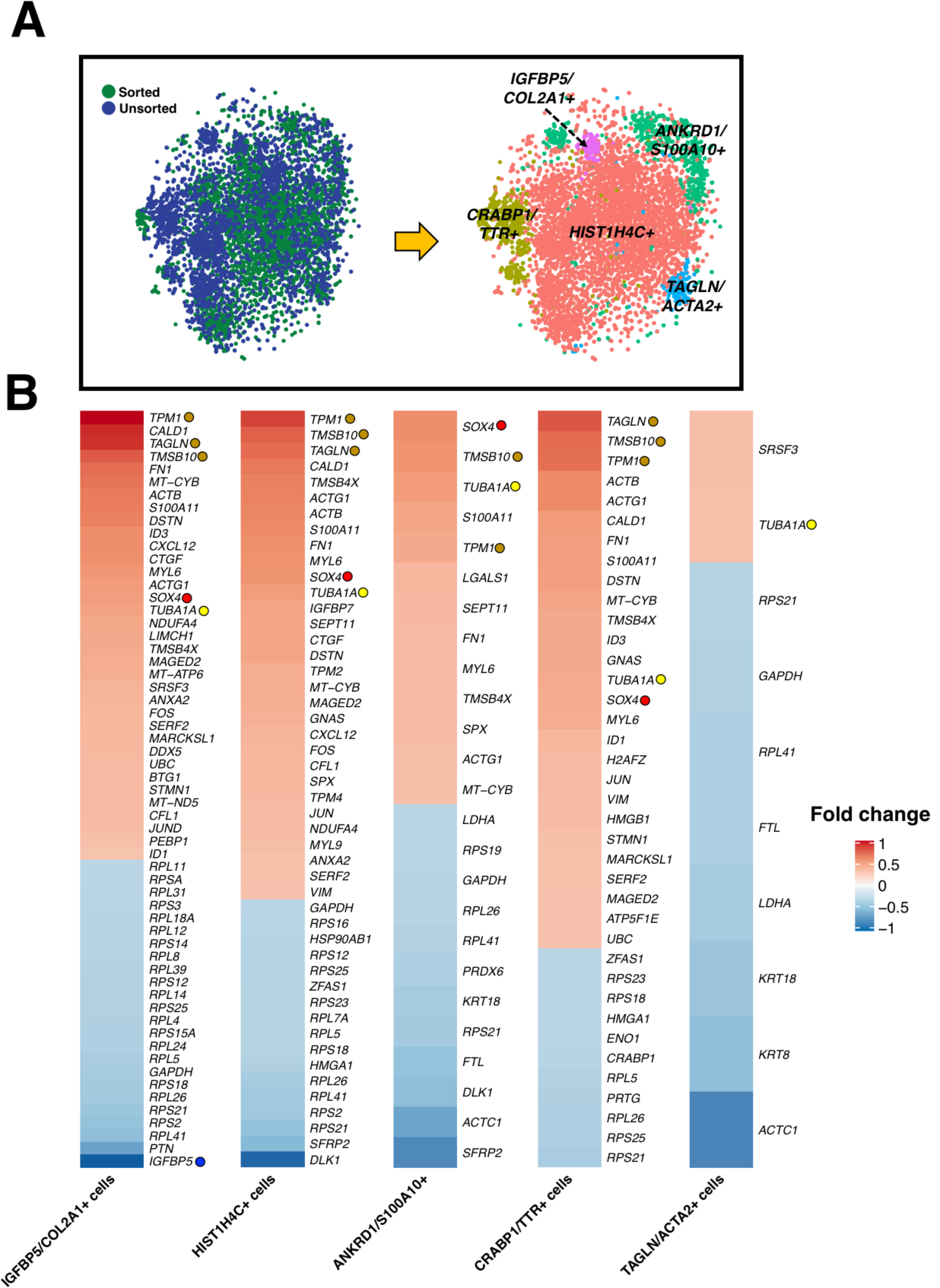
CCA for integrated analysis of sorted and unsorted scRNA-seq datasets. (A) Five major conserved populations were identified after CCA alignment of the sorted and unsorted chondroprogenitor cells (B) DEG analysis indicated that sorted cells exhibited significantly up-regulated expression of several mesenchymal genes including *TPM1, TAGLN* and *TMSB10* (brown circle), which have been suggested to be essential in chondrogenesis. Proliferative markers including *SOX4* (red circle) and *TUBA1A* (yellow circle) were increased, but *IGFBP5* (blue circle) and several ribosomal genes were decreased in sorted cells.

### Sorting improved matrix production and homogeneity in cartilaginous pellets

Sorted and unsorted cells from both the reporter and wildtype lines underwent chondrogenesis in pellet culture for 28 days. Pellets stained with safranin-O for sulfated glycosaminoglycans (sGAGs) showed that sorting increased matrix production as well as homogeneity of cell morphology (**Figure 5A**). Additionally, the layer of non-cartilaginous-like cells surrounding unsorted cell pellets was eliminated in the pellets derived from sorted cells. Biochemical analysis demonstrated that sorting significantly increased the ratio of sGAGs to DNA in pellets by over 20-fold (unsorted: 1.4 ng/ng vs. sorted: 31.8 ng/ng, **Figure 6A**). Similarly, there was an increase in production and homogeneity observed in sorted pellets labeled for COL2A1 (**Figure 5B**). In addition, IHC labeling for COL1A1 showed a slight decrease while that labeling for COL10A1 showed an increase in the respective matrix proteins with sorting (**Figure 5C and 5D**). Additionally, pellets formed with sorted cells had more concentrated staining of COL6A1 around the cells as shown with IHC compared to the more diffused pattern observed with unsorted cells (**Figure S4**).

**Figure 5.**
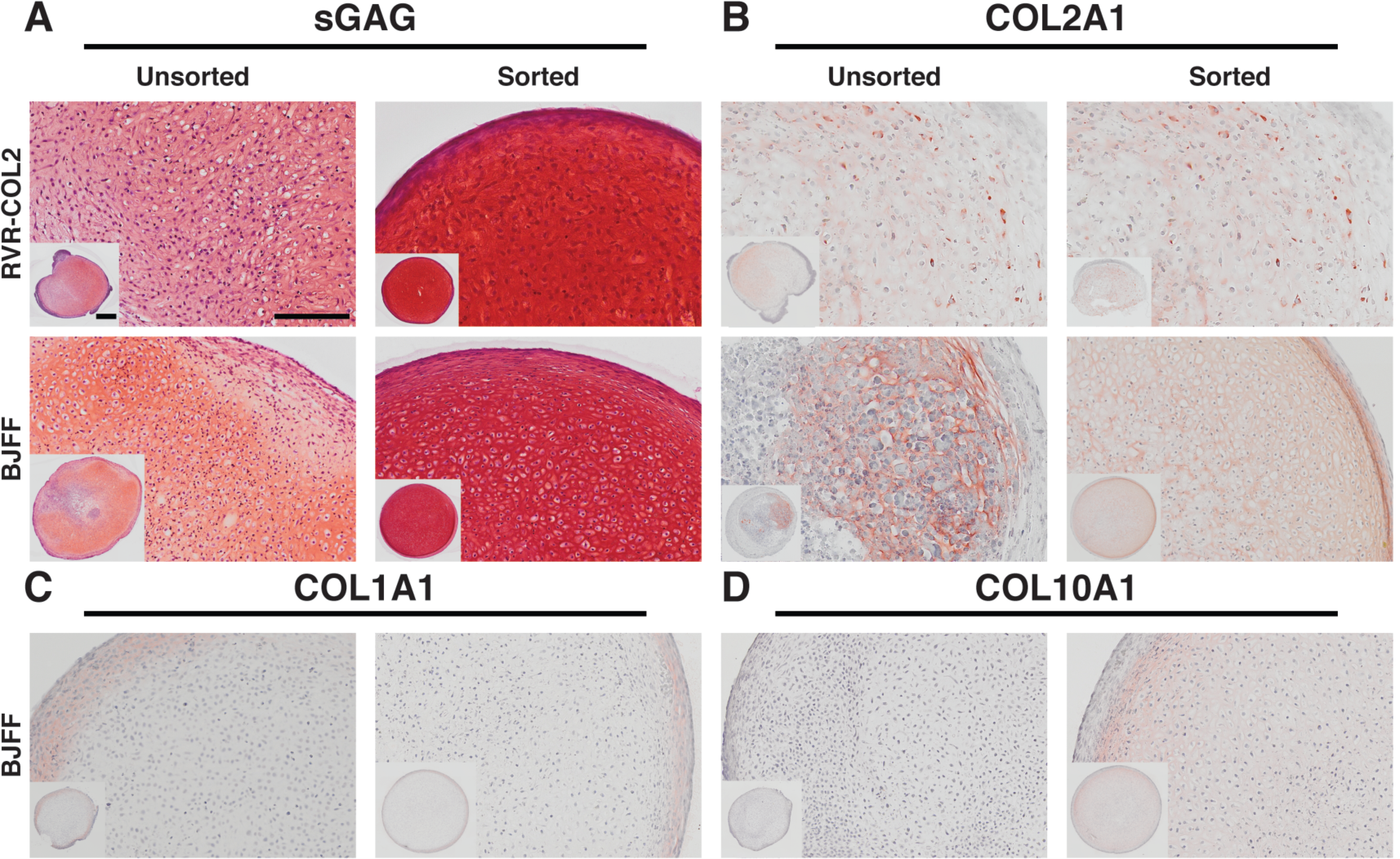
Histology and IHC for matrix proteins in RVR-COL2 and BJFF pellets. (A) Safranin-O staining for sGAG showing pellets derived from sorted chondroprogenitor cells had more robust staining and homogenous cell morphology compared to pellets derived from unsorted cells in both lines. (B) Labeling of COL2A1 showed similar results with an increase in COL2A1 in sorted pellets as opposed to unsorted which has isolated areas of staining. (C) There was little labeling of COL1A1 for both unsorted and sorted cell pellets. (D) Labeling for COL10A1 was increased with sorting. See also Figure S4.

**Figure 6.**
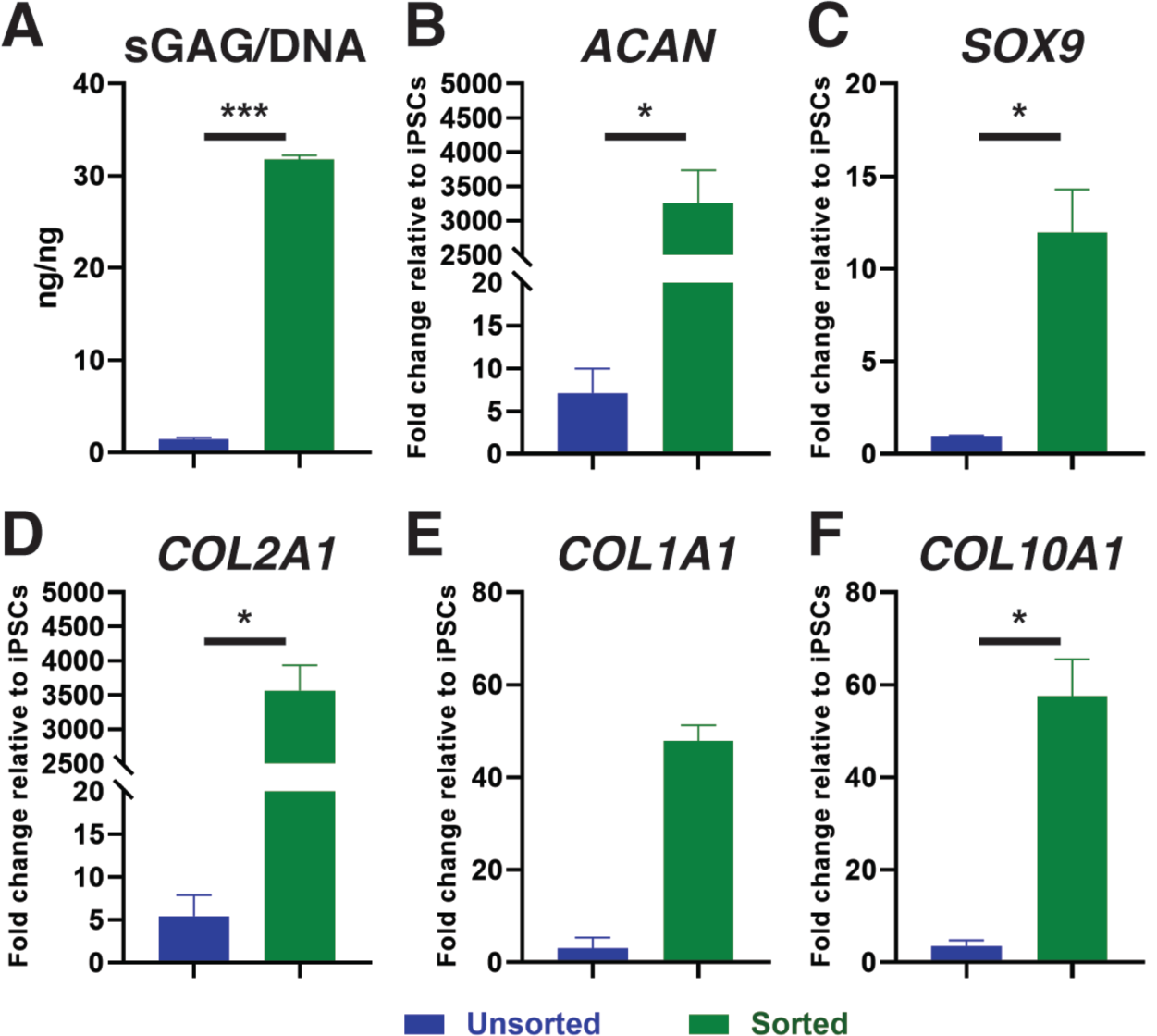
Quantitative analysis of matrix production and gene expression. (A) Sorting of chondroprogenitor cells prior to chondrogenesis significantly increased the sGAG/DNA ratio to over 30 ng/ng. (B-D) Expression of chondrogenic genes *ACAN, SOX9,* and *COL2A1* was significantly increased with sorting. (E) Fibrocartilage and bone matrix marker, *COL1A1*, trended towards an increase with sorting. (F) Sorting significantly unregulated hypertrophic cartilage marker *COL10A1*. Gene expression in reference to undifferentiated hiPSCs with housekeeping gene TBP. * p < 0.05. *** p < 0.001. Data represented as mean ± SEM. n = 3-4 pellets (technical replicates)/group.

### Expression of cartilaginous genes was significantly higher in pellets derived from triple positive chondroprogenitor cells

Gene expression in pellets derived from unsorted and triple positive-sorted chondroprogenitor cells was analyzed using RT-qPCR. Chondrogenic genes *SOX9* (unsorted: 0.98-fold change vs sorted: 11.97 fold change), *ACAN* (unsorted: 7.11-fold change vs. sorted: 3254-fold change), and *COL2A1* (unsorted: 5.42-fold change vs. sorted: 3566-fold change) were significantly increased in sorted pellets (**Figure 6B-D)**. Additionally, *COL1A1* trended toward significance (p = 0.0571) (unsorted: 3.11-fold change vs. sorted: 47.85-fold change) and *COL10A1* was significantly higher in sorted pellets compared to unsorted (unsorted: 3.54-fold change vs. sorted: 57.56-fold change) (**Figure 6E-F**).

### Chondrogenic capacity was maintained through one passage of unsorted and sorted chondroprogenitor cells

Pellets derived from passage 1 (p1) sorted cells exhibited the most robust and homogenous safranin-O staining as compared to the pellets derived from sorted cells of later passages and to the pellets derived from unsorted cells of a similar passage (**Figure S5**). Pellets derived from p2-4 unsorted and sorted chondroprogenitor cells had comparable staining and cell morphology with decreased chondrogenic capacity (**Figure S5)**.

## DISCUSSION

Using a *COL2A1-*GFP reporter line, we have identified a novel combination of surface markers (i.e., PDGFRβ^+^/CD146^+^/CD166^+^/CD45^-^) depicting a unique progenitor population with robust chondrogenic potential in hiPSC chondrogenesis. This finding was further confirmed by significantly increased cartilaginous matrix production of the prospectively isolated cells with these selected markers from a wildtype, non-edited hiPSC line. The results of scRNA-seq of sorted cells revealed that cells positive for PDGFRβ, CD146, and CD166 exhibited enhanced cell homogeneity with decreased neurogenic subpopulations. These findings support the hypothesis that sorting of hiPSC-derived chondroprogenitor cells using surface markers can be used to purify progenitor cells with enhanced chondrogenic potential, without the need for genetic modification to improve hiPSC chondrogenesis (Adkar et al., 2018; Diekman et al., 2012).

We previously reported that chondroprogenitor cells at the end of mesodermal lineage differentiation had high expression of CD146 and CD166 (Adkar et al., 2018). In the present study, we observed that these markers were also highly co-expressed with *COL2A1*. CD146 and CD166, along with CD105, have also been shown to be expressed in chondroprogenitors in articular cartilage (Alsalameh et al., 2004; Su et al., 2015; Vinod et al., 2018). While our chondroprogenitor cells did not co-express CD105 (ENG) with *COL2A1,* sorting did enrich *CD105* expression. Interestingly, it has been shown that CD105 itself may not indicate chondrogenic potential (Cleary et al., 2016). In addition, scRNA-seq showed that sorted cells exhibited increased expression of *ITGB1* (*CD29*) and *ITGA5* (*CD49e*), which have been deemed necessary for chondrogenic differentiation in progenitor cells and MSCs (Cicione et al., 2010; Vinod et al., 2018; Williams et al., 2010). Nevertheless, our chondroprogenitor cells had somewhat different expression profiles than skeletal progenitor cells identified previously *in vivo* (Chan et al., 2018; Wu et al., 2013). Moderate expression of CD164, a surface marker of the skeletal stem cell (Chan et al., 2018), was conserved between the unsorted and sorted chondroprogenitor cells while many other markers described were absent from both populations including prechondrocyte markers BMPR1β and CD73 (NT5E) (Wu et al., 2013). Therefore, the chondroprogenitor population described in this study is a distinct, unique subpopulation of iPSCs that possesses robust chondrogenic potential.

Several factors may contribute to the differences in cell surface markers that have identified as markers of chondrogenesis in these different cell types. First, in our study we used a differentiation protocol which follows the paraxial mesodermal lineage of cartilage (Adkar et al., 2018; Loh et al., 2016). Different types of cartilage follow various developmental pathways (e.g., paraxial mesoderm vs. lateral plate mesoderm) and therefore the other studies could be investigating these lineages, thus the cells would have different surface marker expression during differentiation (Bronner and Ledouarin, 2012; Decker et al., 2014; Loh et al., 2016). Another explanation may be the time point along the developmental pathway in which the cells are being investigated. Our surface marker profiles are based on the expression of *COL2A1*. While COL2A1 is one of the most prominent matrix proteins in articular cartilage (Sophia Fox et al., 2009) and can indicate chondrogenic potential and determination of a chondrogenic fate (Grant et al., 2000), *COL2A1* is a relatively late marker of chondrogenesis (Kosher and Solursh, 1989). Therefore, differences between the cell surface markers identified in our study as compared to other previous work may reflect differences in the prescribed differentiation pathway or the specific subpopulation identified.

In addition to the fact that *COL2A1* expression is a later chondrogenic marker, *COL2A1* expression was found throughout the entire unsorted populations including neurogenic cells, indicating that *COL2A1*+ cells were heterogenous. This finding is consistent with studies showing that *COL2A1* expression may be a broader indicator for the initial lineage specification of a variety of tissues rather than a sole marker for chondrogenesis during embryonic development (Kosher and Solursh, 1989; Kulyk et al., 1991; Nah et al., 1988). Indeed, it has been reported that *COL2A1* is expressed in the floor plate of the central nervous system (Yan et al., 1995), which provides a plausible explanation for our observation of *COL2A1* expression in neurogenic cells. Following sorting, the size of the chondrogenic *SOX9/COL2A1* population was increased and, while the neural *SOX2* populations were reduced, a *SOX2/TTR* population still remained. In fact, this population had high expression of CD47, an integrin-associated and modulating protein (Brown and Frazier, 2001) that could be used as an additional marker for sorting in future experiments to improve homogeneity. The expression of nestin and several mesenchyme markers appeared to be permissive in sorted cells, suggesting that PDGFRβ/CD146/CD166 triple-positive cells may have a similar signature as neural crest cells(Cile Coste et al., 2017; Isern et al., 2014) and might come primarily from mesenchyme populations in unsorted cells. Nonetheless, despite the presence of 6 unique cell clusters, including the *SOX2/TTR* population, sorted chondroprogenitor cells showed robust chondrogenic capacity.

The sorted chondroprogenitors, which all express PDGFRβ, CD146, and CD166, were found to be localized in the mesenchyme clusters of unsorted cells. Alignment of the unsorted and sorted populations by CCA allowed us to compare similarities and differences between the two groups. After alignment, the largest cell cluster expressed histone H4 (*HIST1H4C*). Histones are primarily synthesized during the S-phase of the cell cycle to package the replicated DNA (Osley, 1991), thus indicating the large portion of cells in both sorted and unsorted populations are proliferative. Since scRNA-seq only captures a snap shot in time of the cells’ RNA profiles, it is difficult to tell which proliferating cells may give rise to the desired populations when they transition to the G1 phase. Additionally, it has been shown that transcriptional activity is significantly altered between the G1 and S/G2/M phases (Zopf et al., 2013). Therefore, it is likely that if the cell cycles were synchronized, there would be more homogenous transcriptional profiles. Furthermore, there was a decrease in insulin-like growth factor binding protein-5 (*IGFBP5)* expression in sorted cells among the *IGFBP5/CO2A1* population compared to unsorted. IGFBP5 plays a role in insulin-like growth factor-1 (IGF-1)-dependent chondrocyte proliferation (Kiepe et al., 2001) and protects cartilage during OA-induced degeneration (Clemmons et al., 2002). This may imply that sorted cells may be precursors not fully committed into chondrogenic linage in comparison with unsorted cells. This could be further supported by the observation that sorted cells had increased expression in neural crest and proliferation markers (i.e., *SOX4* and *TUBA1A*, respectively) (Sviderskaya et al., 2009). Indeed, for all populations identified in the sorted cells, we found that they exhibited elevated expression in proliferative and mesenchymal genes, further suggesting that sorted cells were primarily derived from mesenchyme populations in unsorted cells. Nonetheless, subpopulations in sorted cells still expressed unique gene signatures as shown by the clustering. This finding implies that chondrocytes may differentiate from mesenchyme cells with a variety of transcriptomic profiles if given the correct signaling cues with appropriate timing.

Cartilaginous pellets derived from sorted chondroprogenitor cells showed a significant increase in chondrogenic matrix production and gene expression along with the elimination of a surrounding layer of non-chondrocyte-like cells. Surprisingly, there was also a relatively small increase in IHC labeling of COL10A1, a matrix protein often associated with hypertrophic chondrocytes (Kronenberg, 2003; Pacifici et al., 1990). Interestingly, COL6A1 was observed to be more localized around the cells in pellets derived from sorted cells. In developing neonatal cartilage, COL6A1 is found throughout the matrix, but with maturity it is only found in the pericellular matrix surrounding the chondrocytes (Guilak et al., 2018; Morrison et al., 1996; Sherwin et al., 1999). The increased expression in COL10A1 at both mRNA and protein levels alongside the co-localization of COL6A1 around chondrocytes suggests that the chondrocytes derived from the sorted cells were at more mature stages as compared to the chondrocytes derived from unsorted cells after 28 days of chondrogenic culture.

As cell sorting can significantly reduce the number of functional cells (Shields et al., 2015), we also examined the effects of cell expansion on differentiation potential of the sorted cells prior to chondrogenesis. Cells in the first passage following sorting exhibit high chondrogenic potential and s-GAG staining in pellet culture. However, in subsequent passages, cells showed signs of dedifferentiation and loss of chondrogenic capacity, similar to that observed in primary chondrocytes (Darling and Athanasiou, 2005) as well as similarly sorted mouse iPSCs (Diekman et al., 2012). We used expansion media similar to that of MSC expansion media due to similarities of the cells. In the future, various expansion medias could be tested including altering the concentration of bFGF as the growth factor has been shown to maintain multipotency and chondrogenic capacity (Coutu and Galipeau, 2011).

In conclusion, we have identified a unique chondroprogenitor population from hiPSCs which expresses PDGFRβ, CD146, and CD166 and has strong chondrogenic potential. While the population does share some characteristics with previously defined chondroprogenitors and traditionally defined MSCs, it has a distinct profile. The methods and findings in this study will contribute to future cartilage tissue engineering and disease modeling studies to improve understanding and treatment of diseases such as osteoarthritis.

## EXPERIMENTAL PROCEDURES

Experimental methods are briefly summarized. A detailed description is provided in supplemental information.

### hiPSC lines and culture

Two hiPSC lines were used in the current study: RVR *COL2A1-GFP* knock-in line (RVR) and BJFF.6 line (BJFF). Both lines were maintained on vitronectin coated plates (VTN-N; Fisher Scientific, Hampton, NH) with daily medium changes. Cells were passaged at approximately 90% confluency and induced into mesodermal differentiation at 40% confluency.

### Mesodermal differentiation

hiPSCs were induced into mesodermal differentiation in monolayer according to the previously published protocol (Adkar et al., 2018). In brief, cells were fed daily with various cocktails of growth factors and small molecules driving lineage differentiation (anterior primitive streak, paraxial mesoderm, early somite, sclerotome, and chondroprogenitor) in differentiation medium. Upon differentiation into the chondroprogenitor stage, cells were dissociated and were used for cell sorting and chondrogenic differentiation as appropriate.

### Fluorescent activated cell sorting (FACS)

Chondroprogenitor cells were resuspended in FACS Buffer, and stained with various antibodies that are conventionally considered as markers for mesenchymal progenitor cells (**Supplemental Table S1**). Cells were then sorted by Aria-II FACS machine.

### 10X Chromium Platform scRNA-seq

Cells were thawed at 37°C and resuspended in PBS + 0.01% BSA at concentration of 2,000 cells/μl. Cell suspension were submitted to the Genome Technology Access Center (GTAC sequencing core) at Washington University in St. Louis for library preparation and sequencing. In brief, 10,000 cells per sample were loaded on a Chromium Controller (10x Genomics) for single capture. Detailed methods for quality control and processing for scRNA-sea data are described in Supplemental Information.

### Expansion of chondroprogenitor cells

Sorted and unsorted chondroprogenitor cells were plated on non-coated flasks and cultured in MEM alpha media (Gibco) with 1% P/S, 50 μg/ml ascorbate, and 10 ng/ml FGF. Cells were fed every three days until 80-90% confluency prior to further expansion or chondrogenesis. chondroprogenitor cells were passaged up to four times.

### Chondrogenic differentiation

Sorted, unsorted, and expanded chondroprogenitor cells were re-suspended at 2.5 × 10^5^ cells / mL in chondrogenic medium supplemented with 10 ng/ml human transforming growth factor beta 3 (TGF-β3; R&D Systems, Minneapolis, MN). Chondrogenic pellets were cultured at 37°C for 28 days. Medium was changed every 3-4 days.

### Histology and Immunohistochemistry

After chondrogenic differentiation, pellets were fixed and sectioned at 8 µm. Slides were either stained with Safranin-O and hematoxylin for glycosaminoglycans evaluation or labeled against various collagen antibodies including COL1A1, COL2A1, COL6A1, and COL10A1.

### Biochemical analysis

Chondrogenic pellets digested. The PicoGreen (Invitrogen, Carlsbad, CA) and dimethymethlyene blue, with chondroitin-4-sulfate as a standard, assays were used according to the protocols to quantify DNA and sGAG respectively.

### Gene expression

Day 28 pellets were homogenized. Gene expression was analyzed using the *ΔΔ*C_T_ method relative to undifferentiated hiPSCs with the reference gene TATA-box-binding protein (*TBP*) (Livak and Schmittgen, 2001). Sequences of primers can be found in the **Supplemental Table S2.**

### Statistical analysis

Gene expression and GAG/DNA data were tested for normality using the Shapiro-Wilk test. An unpaired t-test with Welch’s correction was then performed assuming a Gaussian distribution. If data was not normal, an unpaired Kolmogorov-Smirnov test was performed. All calculations were performed using GraphPad Prism software. Sorting experiments were run 7 separate times with technical replicates of 3-4 each time. Two-tailed p values were calculated and reported at a 95% confidence interval.

## AUTHOR CONTRIBUTIONS

Conceptualization: A.D., C.L.W., F.G.; Methodology: A.D., C.L.W., S.S.A.; Investigation, A.D., C.L.W., N.S.; Writing – Original Draft, A.D., C.L.W., F.G.; Writing – Review and Editing, A.D., C.L.W., N.S., S.S.A., C.A.G., F.G.; Supervision, C.A.G., F.G.; Funding Acquisition, C.A.G., F.G.

## ACKNOWLEDGMENTS

This work was supported by the Nancy Taylor Foundation, Arthritis Foundation, NIH (AG46927, AG15768, AR67467, AR65956, T32 DK108742, T32 EB018266), NSF EAGER Award, and Taiwan GSSA Scholarship. The authors would like to thank Christopher Sawyer from the Genome Technology Access Center (GTAC) and Erica Lantelme, Dorjan Brinja, and Ananya Mitra from the Flow Cytometry & Fluorescence Activated Cell Sorting Core, Washington University in St. Louis for their assistance.

