## Supplemental Information for "Prospective Isolation of Chondroprogenitors from Human iPSCs Based on Cell Surface Markers Identified using a CRISPR-Cas9-Generated Reporter"

**SUPPLEMENTAL FIGURES AND LEGENDS**

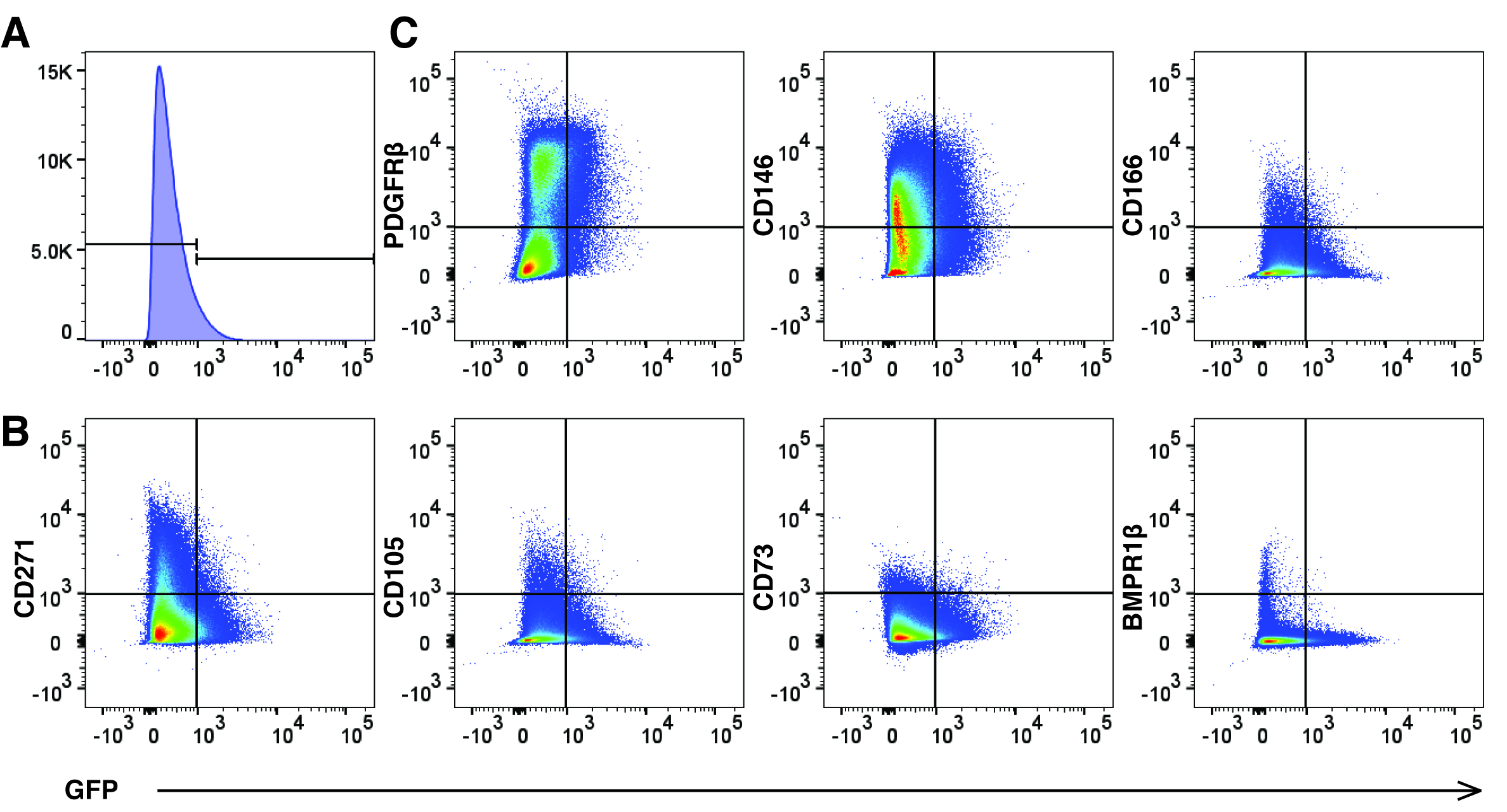

**Figure S1 related to Figure 1.** **Representative flow plots for surface markers co-expressing *COL2A1-GFP* in RVR chondroprogenitor cells.** (A) Approximately 4.27% of cells expressed *COL2A1-GFP*. (B) Most *COL2A1*-GFP+ cells did not express CD271, CD105, CD73, and BMPR1β. (C) PDGFRβ, CD146, and CD166 were co-expressed with *COL2A1*-*GFP*.

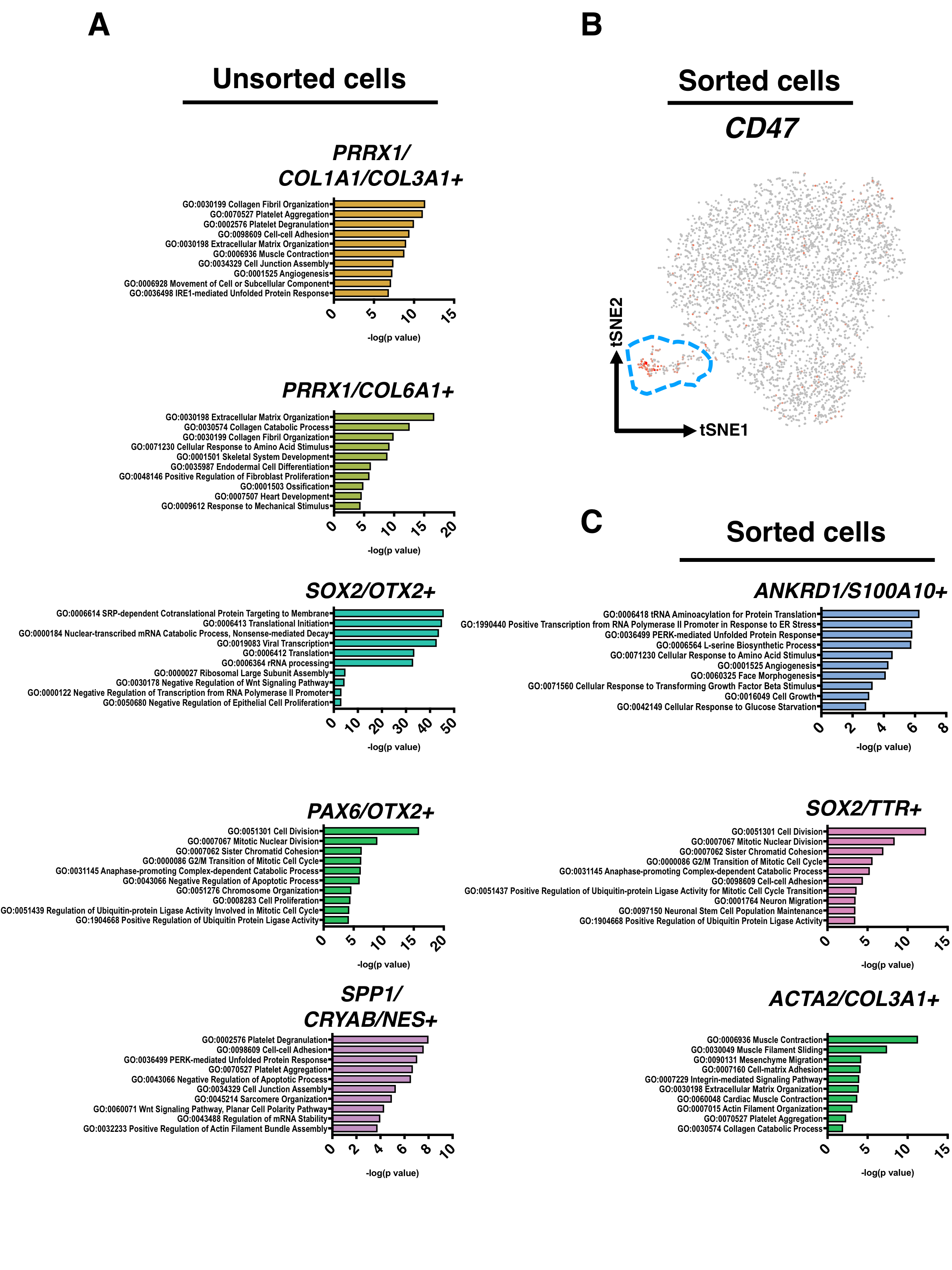

**Figure S2 related to Figure 2 and 3. GO enrichment analysis of unsorted and sorted cells.** (A) Top 10 GO terms (biological process) that were associated with each population in unsorted cells. (B) CD47 was highly expressed in *SOX2/TTR+* cells. (C) Top 10 GO terms (biological process) that were associated with each population in sorted cells.

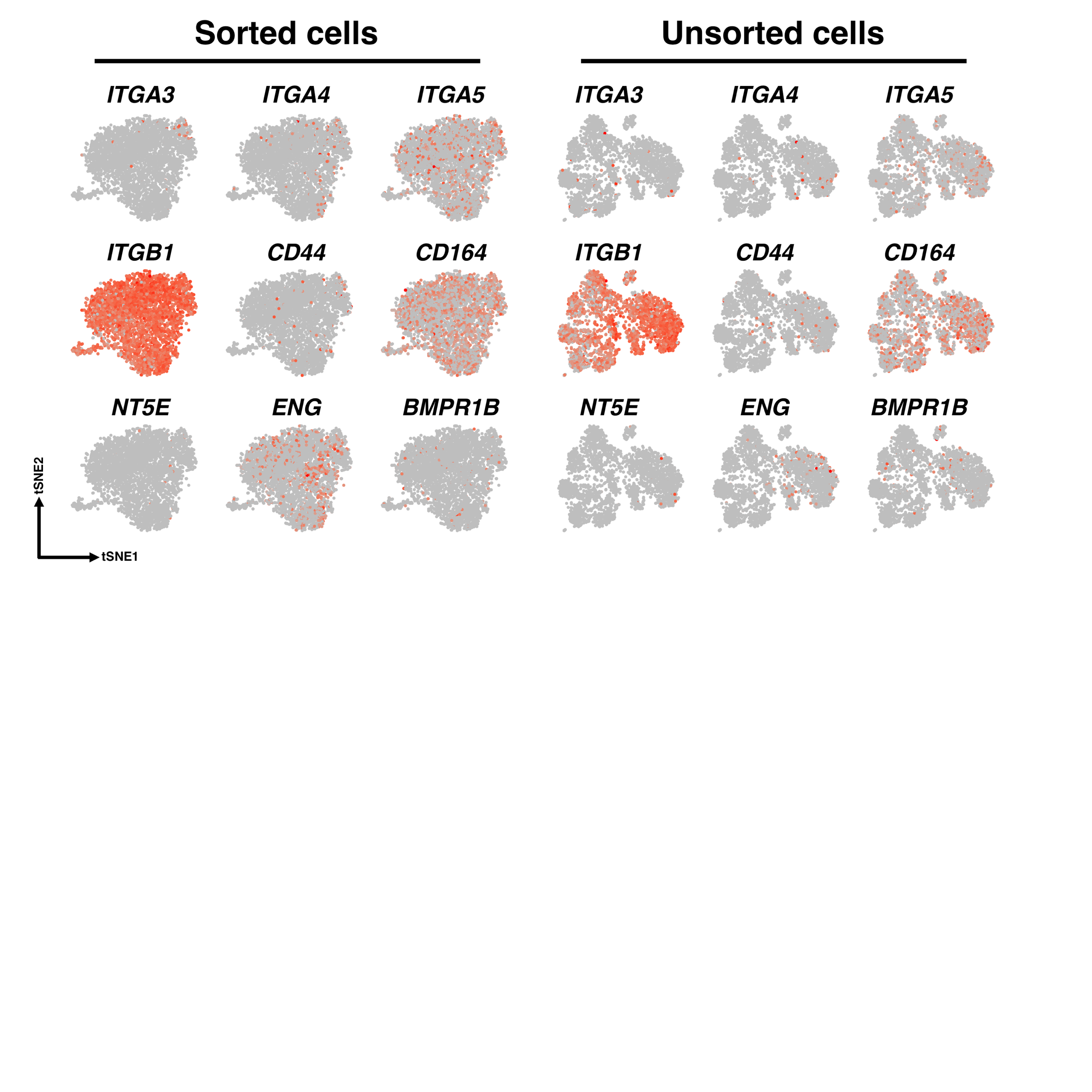

**Figure S3 related to Figure 2 and 3. Expression profiles of pro-chondrogenic genes in sorted and unsorted chondroprogenitor cells.** scRNA-seq reveals that sorted and unsorted cells had distinct gene expression patterns of several markers that were proposed to be pro-chondrogenic identified by previous studies.

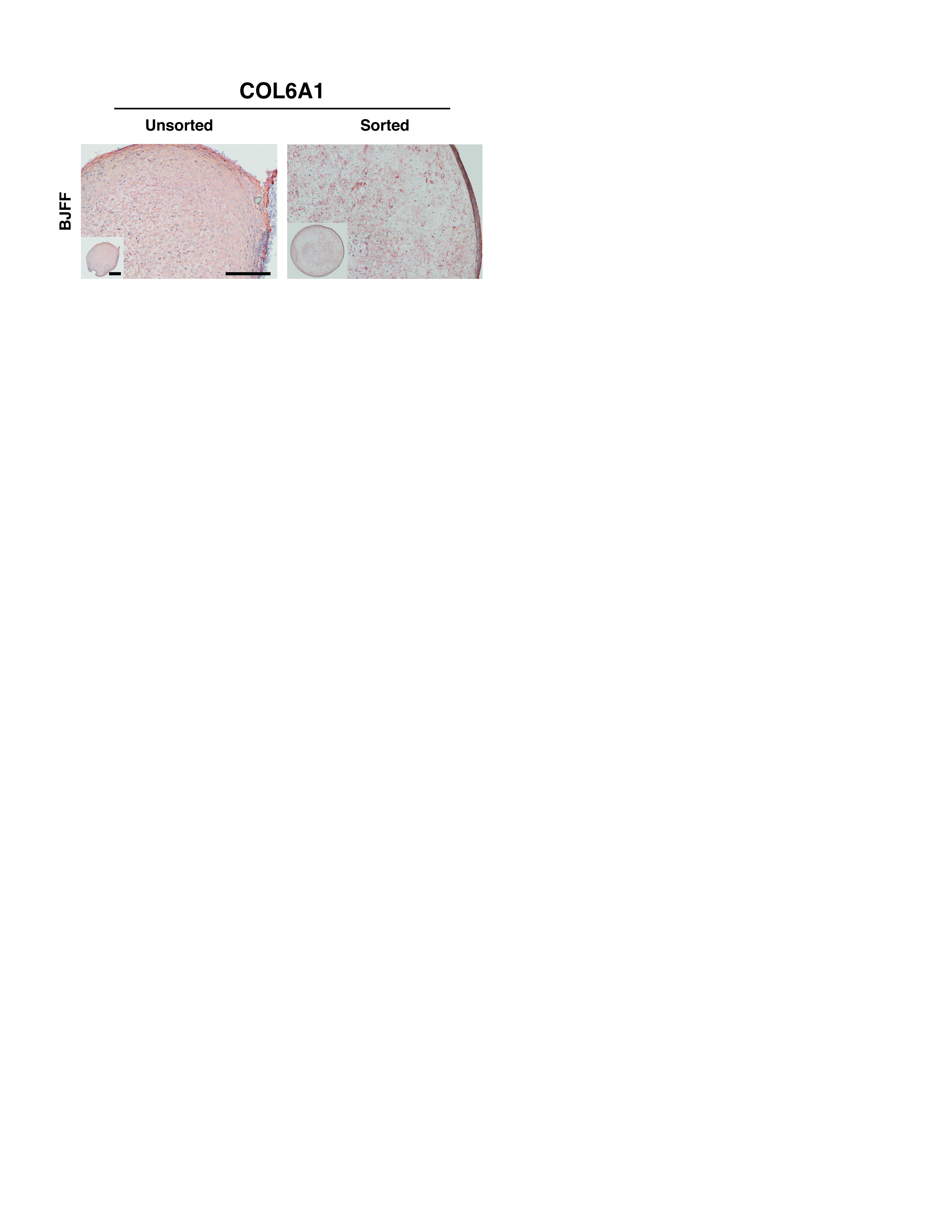

**Figure S4 related to Figure 5. IHC labeling for COL6A1**. There was more distributed labeling for COL6A1 in unsorted chondroprogenitor pellets compared to the localization around cells in sorted chondroprogenitor pellets.

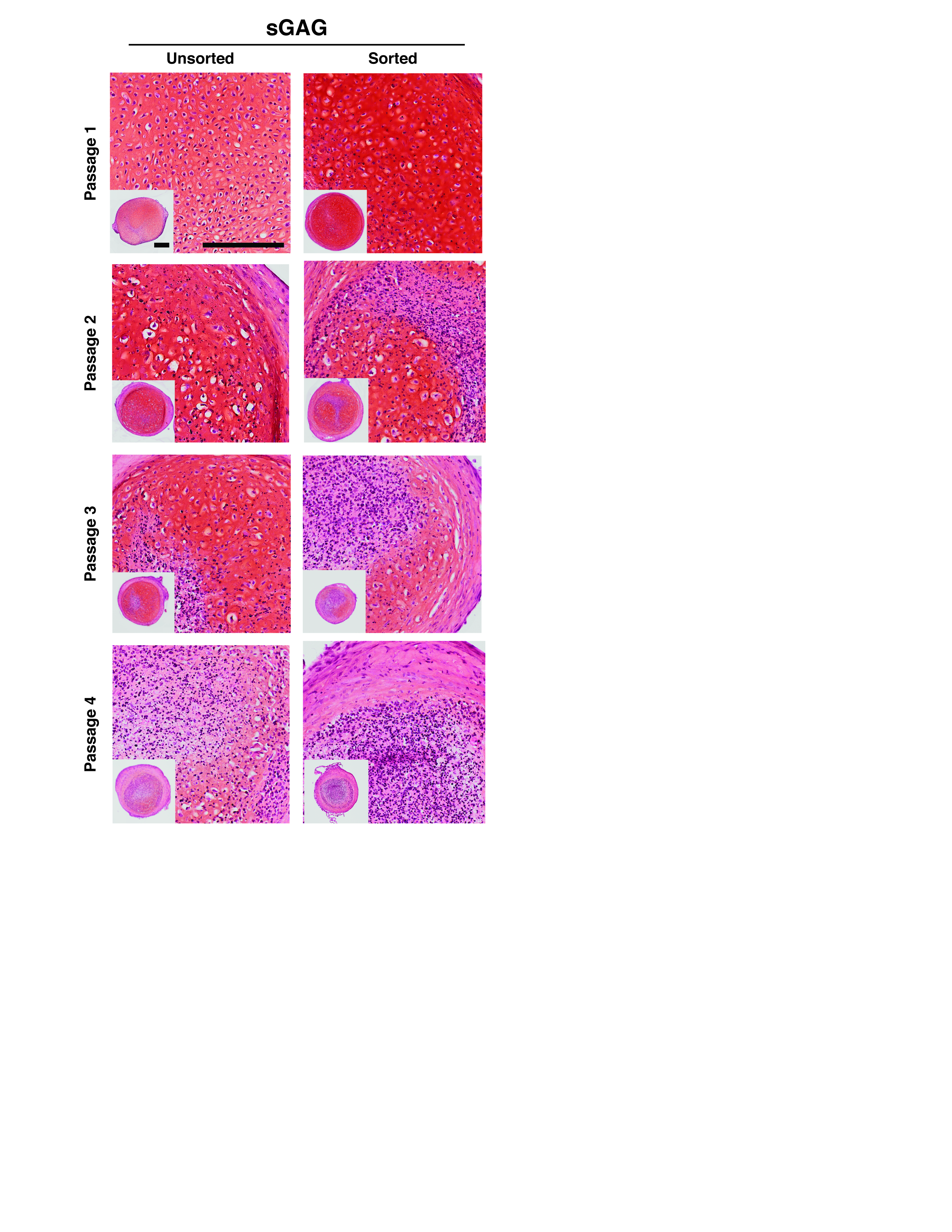

**Figure S5. Histology of pellets derived from *in vitro* expanded unsorted and sorted chondroprogenitors.** Chondrogenic capacity was maintained after one passage of both unsorted and sorted chondroprogenitor cells as shown by staining for sGAG. There was more robust staining in pellets derived from sorted cells. Safranin-O staining for sGAG showed similar loss of chondrogenic capacity for both unsorted and sorted chondroprogenitor cells through four passages.

**Table S1. Antibodies used for flow cytometry and sorting.**

| **Marker** | **Color** | **Company** | **Catalog Number** | |
| --- | --- | --- | --- | --- |
|  |  |  | **Antibody** | **Isotype** |
| CD45 | FITC | BioLegend | 304006 | 400110 |
| CD166 | PE | BioLegend | 343904 | 400112 |
| CD146 | PE/Cy7 | BioLegend | 361008 | 400126 |
| PDGFRβ | APC | BioLegend | 323608 | 400121 |
| CD271 | PE/Cy7 | BioLegend | 345110 | 400126 |
| CD105 | PE | BioLegend | 323206 | 400113 |
| CD73 | Pacific Blue | BioLegend | 244012 | 400151 |
| BMPR1β | APC | R&D Systems | FAB5051A | IC0041A |

**Table S2.** **Human primer sequences.** Primers were used for RT-qPCR and are listed as 5’ to 3’.

| **Gene** | **Forward Sequence** | **Reverse Sequence** |
| --- | --- | --- |
| *ACAN* | CACTTCTGAGTTCGTGGAGG | ACTGGACTCAAAAAGCTGGG |
| *COL1A1* | TGTTCAGCTTTGTGGACCTC | TTCTGTACGCAGGTGATTGG |
| *COL2A1* | GGCAATAGCAGGTTCACGTA | CTCGATAACAGTCTTGCCCC |
| *COL10A1* | CATAAAAGGCCCACTACCCAAC | ACCTTGCTCTCCTCTTACTGC |
| *SOX9* | CGTCAACGGCTCCAGCAAGAACAA | GCCGCTTCTCGCTCTCGTTCAGAAGT |
| *TBP* | AACCACGGCACTGATTTTCA | ACAGCTCCCCACCATATTCT |

**SUPPLEMENTAL EXPERIMENTAL PROCEDURES**

Contact for Reagents and Resource Sharing. Further information and requests for resources and reagents should be directed to and will be fulfilled by the Lead Contact, Farshid Guilak.

**Experimental Model and Subject Details**

**hiPSC lines and culture.**

Two hiPSC lines were used in the current study: RVR *COL2A1-GFP* knock-in line (RVR) and BJFF.6 line (BJFF). The RVR line was retrovirally reprogrammed from BJ fibroblasts and characterized as previously described (Lee et al., 2012, 2015). Using CRISPR-Cas9 genome editing, a GFP reporter allele was added to the *COL2A1* gene as previously reported (Adkar et al., 2018). The cells were cultured in mTeSR1 medium (StemCell Technologies, Vancouver, CA). The BJFF line was derived using Sendai viral transfection of foreskin fibroblasts from a newborn male and cultured in Essential 8 Flex medium (E8; Life Technologies, Carlsbad, CA). Both lines were maintained on vitronectin coated plates (VTN-N; Fisher Scientific, Hampton, NH) with daily medium changes. Cells were passaged at approximately 90% confluency and induced into mesodermal differentiation at 40% confluency.

**Mesodermal differentiation.**

hiPSCs were induced into mesodermal differentiation in monolayer according to the previously published protocol (Adkar et al., 2018). In brief, cells were fed daily with various cocktails of growth factors and small molecules driving stepwise lineage differentiation (anterior primitive streak, paraxial mesoderm, early somite, sclerotome, and chondroprogenitor) in differentiation medium composed of IMDM GlutaMAX (IMDM; Fisher Scientific, Waltam, MA) and Ham’s F12 Nutrient Mix (F12; Fisher Scientific) with 1% chemically defined lipid concentrate (Gibco, Gaithersberg, MD), 1% insulin/human transferrin/selenous acid (ITS+; Corning, Durham, NC), 1% penicillin/streptomyocin (P/S; Gibco), and 450 μM 1-thioglycerol (Sigma–Aldrich, St. Louis, MO). Upon differentiation into the chondroprogenitor stage, cells were dissociated using TrypLE (Gibco) at 37ºC followed by neutralization with equal parts of DMEM/F-12, GlutaMAX^TM^ (DMEM/F12; Gibco) with 10% fetal bovine serum (FBS; Atlanta Biologicals, Flowery Branch, GA) and 1% P/S. The dissociated cells were used for cell sorting and chondrogenic differentiation as appropriate.

**Fluorescent activated cell sorting (FACS).**

Chondroprogenitor cells were resuspended in FACS Buffer (PBS-/- with 1% FBS and 1% penicillin/streptomyocin/fungizone (P/S/F; Gibco) at approximately 40 x 10^6^ / mL. The cells were treated with ~6.5% of Human Tru Stain FC X^TM^ (BioLegend, San Diego, CA) for 10 min at RT. Approximately 5 x 10^4^ cells in 100 µl were used for each compensation. Cells were labeled with appropriate antibodies including an isotype control. A table of the antibodies used can be found in **Supplemental Table S1**. Cells were incubated for 30 minutes at 4ºC and washed with FACS buffer twice. Samples were resuspended in sorting medium consisting of DMEM/F12 with 2% FBS, 2% P/S/F, 2% Hepes (Gibco), and DAPI (BioLegend) at 4 x 10^6^ cells / mL and filtered through a 40 µm cell strainer. Cells were stored on ice prior to sorting.

An Aria-II FACS machine was used to compensate and gate the samples. Sorted cells were collected in collection media composed of DMEM/F12, 20% FBS, 1% P/S/F, 1% ITS+, 1% MEM non-essential amino acids (NEAA; Corning), and 0.1% ROCK inhibitor (Stemcell Technologies). Data was analyzed using FlowJo software.

**10X Chromium Platform scRNA-seq.**

Cells were thawed at 37ºC and resuspended in PBS + 0.01% BSA at concentration of 2,000 cells/μl. Cell suspension were submitted to the Genome Technology Access Center (GTAC sequencing core) at Washington University in St. Louis for library preparation and sequencing. In brief, 10,000 cells per sample were loaded on a Chromium Controller (10x Genomics) for single capture. Libraries were prepared using Single Cell 3′ Library & Gel Bead Kit v2 (#120237 10x Genomics) following manufacture’s instruction. A single cell emulsion (Gel Bead-In-EMulsions, GEMs) is created by making barcoded cDNA unique to each individual emulsion. Recovery agent was added to break GEM and cDNA was then amplified. A library is produced via end repair, dA-tailing, adaptor ligation, post-ligation cleanup with SPRIselect, and sample index PCR. The quality and concentration of the amplified cDNA was evaluated by Bioanalyzer (Agilent 2100) on a High Sensitivity DNA chip (Agilent, #5065-4401). Only cDNA with average library size 260-620 bp were used for sequencing. Sequencing was performed by Illumina HiSeq2500 with the following read length: 26 bp for Read1, 8 bp for i7 Index, and 98 bp for Read2. A species mixing experiment (mouse and human) was also performed prior to running on actual sample to ensure good single cell capture (i.e., cell doublet rate < 5%).

**Preprocessing of scRNA-seq Data.**

Paired-end sequencing reads were processed by Cell Ranger (10x Genomics software, version 2.0.0). Briefly, reads were aligned to the GRCh38 (version 90) for genome annotation, demultiplexing, barcode filtering and gene quantification. Cell Ranger also removes any barcode that has less than 10% of the 99^th^ percentile of total unique molecular identifiers (UMI) counts per barcode as these barcodes are considered to be associated with empty droplets. After this quality control, gene barcode matrix for sorted and unsorted cells were generated by counting the number of UMIs for a given gene (as row) in individual cell (as column). For sorted cell sample, 8,682 cells with 19,140 genes were captured, while for unsorted cell sample, we obtained 8,220 cells with 19,604 genes.

**Unsupervised Clustering Analysis and Cell Cluster Annotation.**

To assess the difference in composition of cell populations due to sorting for selective markers, we performed global unsupervised clustering analysis on data sets of scRNA-seq from sorted and unsorted cells. First, gene barcode matrices from were input into Seurat R package (version 2.4) (Butler et al., 2018). We then removed the low-quality cells with less than 200 or more than 7,000 detected genes, or if their mitochondrial gene content was more than 5%. Genes were filtered out that were detected in less than 3 cells. After filtering out low-quality cells or cell doublets, the gene expression was then natural log transformed and normalized for scaling the sequencing depth to 10,000 molecules per cell. Next, to reduce the variance introduced by “unwanted” sources, we regressed out variation in gene expression driven by cell cycle stages and mitochondrial gene expression with *vars.to.regress* argument in function *ScaleData* in Seurat. We then used *FindVariableGenes* function in Seurat to identify highly variable genes across cells for downstream analysis. These quality control steps resulted in 4,173 cells with 1,833 highly variable genes for sorted cell sample, and 8,630 cells with 3,009 highly variable genes for unsorted cell sample. Dimensionality reduction on the data was then performed by computing the significant principal components on highly variable genes. We then performed unsupervised clustering by using *FindClusters* function in Seurat with the resolution argument set to 0.6, and clusters were then visualized in t-distributed Stochastic Neighbor Embedding (tSNE) plot (van der Maaten and Hinton, 2008).

Differentially expressed genes (DEGs) among each cell cluster were determined using *FindAllMarkers* function in Seurat. DEGs expressing at least in 25% cells within the cluster and with a fold change of more than 0.25 in log scale were considered to be marker genes of the cluster. To determine the biology functions of the marker genes from a given cluster, we performed Gene Ontology (GO) enrichment analysis by using The DAVID Gene Functional Classification Tool (<http://david.abcc.ncifcrf.gov>; version 6.8) (Wei Huang et al., 2007). By comparing these unique biological GO terms with existing RNA-seq datasets and the literature, we were able to annotate cell clusters. Additionally, the top 10 enriched GO terms from biological function category with associated p values were visualized by GraphPad Prism (version 8.0; GraphPad Software).

**Cell Cycle Analysis.**

*CellCycleScoring* function in Seurat was used to determine a cell cycle score on each cell according to its gene expression of G2/M phase (54 genes) and S phase (43 genes) markers (Tirosh et al., 2016). Based on this scoring system, fractions of each cell cluster with given cell cycle score in total cell population were computed and plotted.

**Canonical Correlation Analysis for Integrated Analysis of Sorted and Unsorted scRNA-seq Data.**

To compare cell types and to identify their associated DEGs between sorted and unsorted cells, we applied canonical correlation analysis (CCA), a computational strategy implemented in Seurat for integrated analysis of multiple datasets, on the scRNA-seq datasets from the cells with or without sorting. First, top 1,000 highly variable genes from two datasets were selected. We then use *RunCCA* function to identify common sources of variation between the two datasets and to merge the two objects into a single dataset. We next determined the top principal components of the CCA by examining a saturation in the relationship between the number of principle components and the percentage of the variance explained using the *MetageneBicorPlot* function. By using selected top principal components, we aligned the CCA subspaces with *AlignSubspace* function, which returns a new dimensional reduction matrix allowing for downstream clustering and DEG analyses. DEG analysis was performed on the cells from different datasets but grouped in the same cluster (i.e. conserved cell types between two conditions) after CCA alignment. The methods for cell clustering, identification of conserved cells types and DEGs as well as annotation of cell clusters were similar to the ones mentioned previously. DEGs in each conserved cell types in response to sorting were visualized by ComplexHeatmap R package (Gu et al., 2016).

**Chondrogenic differentiation.**

Sorted, unsorted, and expanded chondroprogenitor cells were re-suspended at 2.5 x 10^5^ cells / mL in chondrogenic medium composed of DMEM/F-12, 1% FBS, 1% ITS+, 55 µM β-mertcaptoethanol, 100 nM dexamethasone (DEX; Sigma-Aldrich), 1% NEAA, 1% P/S, 10 ng/ml human transforming growth factor beta 3 (TGF-β3; R&D Systems, Minneapolis, MN), 50 μg/ml L-ascorbic acid 2-phosphate (ascorbate; Sigma-Aldrich), and 40 μg/ml L-Proline (proline; Sigma-Aldrich). 2 mL of the cell solution was added to a 15mL conical tube and centrifuged for 5 min at 300g to form a pellet. Chondrogenic pellets were cultured at 37ºC for 28 days. Medium was changed every 3-4 days.

**Histology.**

After chondrogenic differentiation, pellets were collected in 10% neutral buffered formalin for fixation for 24 hours. Pellets were then transferred to 70% ethanol, dehydrated, and embedded in paraffin wax. Wax blocks were sectioned at 8 µm and stained for glycosaminoglycans and cell nuclei according to the Safranin-O and hematoxylin standard protocol.

**Immunohistochemistry.**

Histologic sections were washed with xylene three times and rehydrated before labeling against COL1A1, COL2A1, COL6A1, and COL10A1. Antigen retrieval was performed with 0.02% proteinase K for 3 min at 37ºC for COL2A1 and COL6A1 and with pepsin for 5 min at RT for COL1A1 and COL10A1 followed by peroxidase quench then serum blocking for 30 min at RT each. Samples were labeled for 1 hour with the primary antibody (COL1A1: 1:800 Abcam 90395; COL2A1: 1:10 Iowa II-II6B3-s; COL6A1 1:1000 Fitzgerald 70F-CR009X; COL10A1 1:200 Sigma C7974) and for 30 min with the secondary antibody (COL1A1, COL2A1, COL10A1: 1:500 Goat Anti-Mouse Abcam 97021; COL6A1: 1:500 Goat Anti-Rabbit Abcam 6720). Reagent C was then used for enzyme conjugation for 20 min at RT followed by AEC for 2.5 min (COL2A1 and COL6A1) or 2 min (COL1A1 and COL10A1) at RT. Finally, samples were counterstained with hematoxylin to reveal cell nuclei for 45 sec.

**Biochemical analysis.**

Pellets were rinsed with PBS after chondrogenic differentiation and digested at 65ºC overnight in 200 µl of 125 μg/ml papain, 100 mM sodium phosphate, 5 mM EDTA, and 5 mM L-cysteine hydrochloride at 6.5 pH. Samples were stored at -80ºC before thawing to measure double stranded DNA and sulfated glycosaminoglycans (sGAG). The PicoGreen (Invitrogen, Carlsbad, CA) and dimethymethlyene blue, with chondroitin-4-sulfate as a standard, assays were used according to the protocols to quantify DNA and sGAG respectively.

**Gene expression.**

Day 28 pellets were rinsed with PBS and snap frozen in 300 µl of Buffer RL (Norgen Biotek, Thorold, ON, Canada) and stored at -80ºC. Samples were thawed and homogenized with zirconia beads (BioSpec Products, Bartlesville, OK) and a miniature bead beater. RNA was isolated from the samples using the Total RNA Purification Kit according to the manufacture’s protocol (Norgen Biotek). Reverse transcription of the RNA was performed using SuperScript VILO Master Mix (Thermo Fisher) to maintain cDNA. Fast SYBR Green Master Mix (Thermo Fisher) was used for quantitative real time polymerase chain reaction (RT-qPCR) according to manufacturer’s instructions on the QuantStudio 3 (Thermo Fisher). Gene expression was analyzed using the ΔΔC_T_ method relative to undifferentiated hiPSCs with the reference gene TATA-box-binding protein (*TBP*) (Livak and Schmittgen, 2001). Sequences of primers can be found in the **Supplemental Table S2.**
